# DNA ligases of *Prochlorococcus marinus*: an evolutionary exception to the rules of replication

**DOI:** 10.1101/2020.05.11.089284

**Authors:** Erik Hjerde, Ashleigh Maguren, Elizabeth Rzoska-Smith, Bronwyn Kirby, Adele Williamson

**Affiliations:** School of Science, University of Waikato, Hamilton 3240, New Zealand; Department of Chemistry, UiT The Arctic University of Norway, Tromsø, N-9037, Norway

## Abstract

DNA ligases, essential enzymes which re-join the backbone of DNA come in two structurally-distinct isoforms, NAD-dependent and ATP-dependent, which differ in cofactor usage. The present view is that all bacteria exclusively use NAD-dependent DNA ligases for DNA replication, while archaea and eukaryotes use ATP-dependent DNA ligases. Some bacteria also possess auxiliary ATP-dependent DNA ligases; however, these are only employed for specialist DNA repair processes. Here we show that in the genomes of high-light strains of the marine cyanobacterium *Prochlorococcocus marinus*, an ATP-dependent DNA ligase has replaced the NAD-dependent form, overturning the present paradigm of a clear evolutionary split in ligase usage. Genes encoding partial NAD-dependent DNA ligases are found on mobile regions in highlight genomes and lack domains required for catalytic function. This constitutes the first reported example of a bacterium that relies on an ATP-dependent DNA ligase for DNA replication and recommends *P. marinus* as a model to investigate the evolutionary origins of these essential DNA-processing enzymes.

## Introduction

DNA ligases, enzymes that join breaks in the phosphodiester backbone of double-stranded DNA, are essential for DNA replication and repair in all organisms. They are classified as ATP-dependent (AD-ligases) or NAD-dependent (ND-ligases) on the basis of the adenylating cofactor used during catalysis (1). AD- and ND-ligases have different taxonomic distributions among cellular organisms and possess distinct sequence and structural features. ND-ligases are almost entirely limited to eubacteria where they are essential for replication, whereas eukaryotes and archaea use AD-ligases. This key difference in cellular replication machinery represents a central delineation between bacterial and archaeal/eukaryotic cell lineages (2); however the reason why bacteria preferentially use of NAD-ligases remains a long-standing question in the field (3). Previously, we described how strains of the marine cyanobacterium *Prochlorococcus marinus* encode multiple AD-ligases in their genomes, yet lack the other subunits associated with known AD-ligase-dependent repair pathways (4). *P. marinus* ecotypes are broadly classified as high-light which reside in the UV-damaging nutrient-poor upper ocean and low-light which experience less UV, and have higher nutrient access (5). Low-light strains have the smallest genomes of any free-living organisms (1.66 −1.75 MB) yet despite this minimization, they encode up to three predicted AD-ligases with different domain compositions (Fig 1A). To gain further insight, we have used a comparative genomics approach to survey the diversity of DNA ligases among sequenced isolates of *P. marinus*, which indicates a role for these AD-ligases in DNA replication.

**Figure 1.**
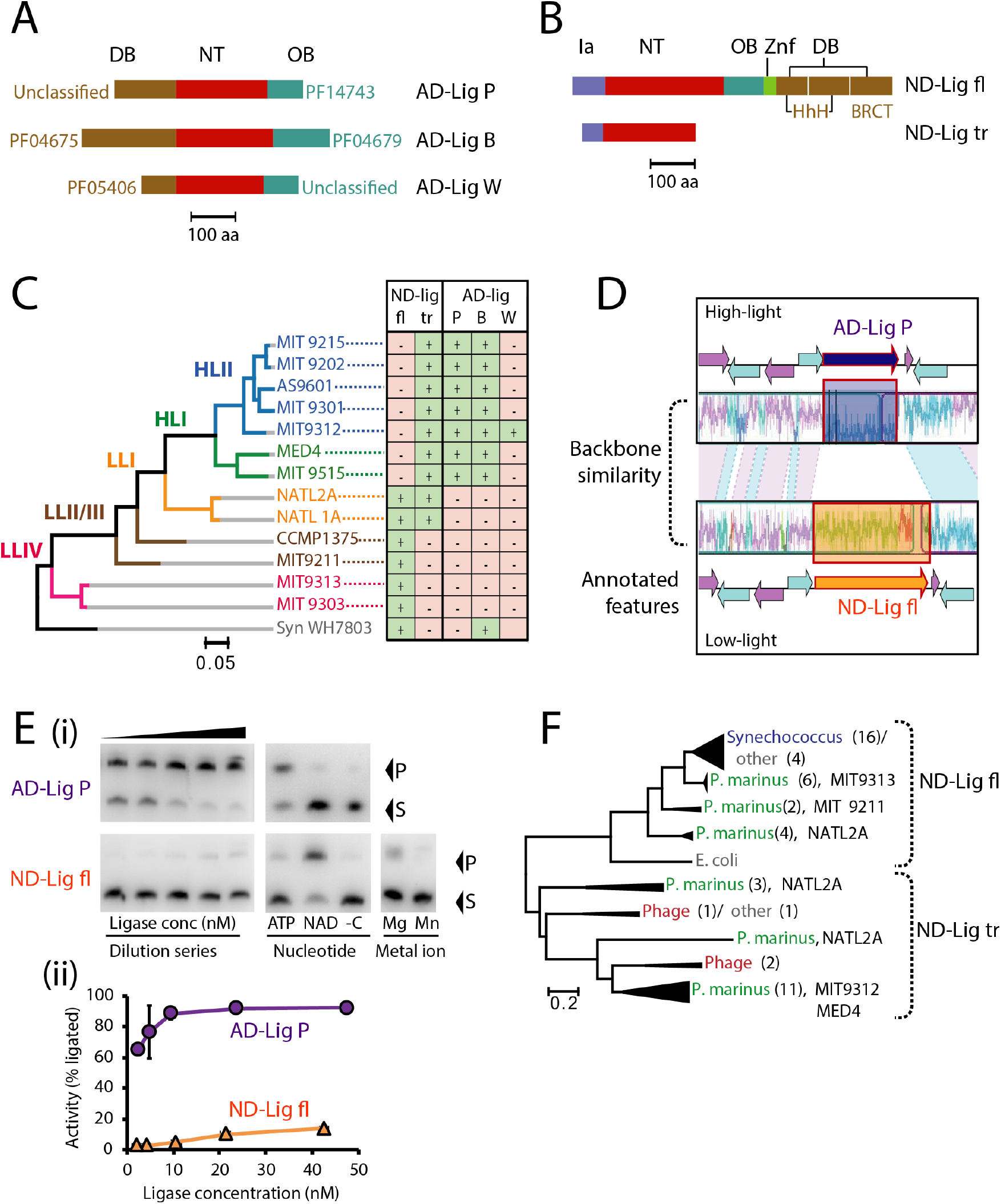
High-light and low-light *P.marinus* ecotypes have different complements of DNA ligases. A) Schematic of ATP-dependent DNA ligases found in high-light *P. marinus*, classified by Pfam assignment as described in (4). Domains are colored by function and Pfam identifiers are given adjacent. The structurally-conserved conserved catalytic core nucleotidyl transferase domain (NT) and oligonucleotide-binding (OB) domains are colored red and cyan; the variable DNA-binding domain (DB) is beige. B) Schematic of full-length (ND-Lig fl) and truncated (ND-lig tr) NAD-dependent DNA ligases colored by domain function. The OB, zinc-finger (ZnF) and DB domains are essential for activity in homologs (6). C) Presence and absence of ATP- and NAD-dependent DNA ligases in the 13 closed genomes of *P. Marinus strains* with *Synechococcus* WH7803 as an outgroup. Data for all 41 closed and genomes and scaffolds is given in data S1. Strain evolution is inferred from the DNA gyrase B gene as described (15), and ecotypes are colored according to their assignment in ProPortal (11) with abbreviations high-light (HL) and low-light (LL). D) Genetic context of the ATP-dependent DNA ligase AD-Lig P and full-length NAD-dependent DNA ligase (ND-Lig fl) in high-light strain MIT9312 and low-light strain NATl2A respectively. Aligned regions of backbone similarity are colored violet/cyan and annotated genes are shown as arrows. No similarity was detected for the region encoding the ligase sequences AD-Lig P and ND-Lig fl. For entire alignment with representatives of all clades see data S2. E) DNA ligase activity for high-light AD-Lig P (MIT9302) and low-light ND-Lig fl (MIT9211). (i) Representative urea PAGE gels of ligation including specific activity by dilution (left), nucleotide cofactor preference (center) and metal ion preference (right). (ii) Ligation as a function of enzyme concentration integrated from left-most panel above. Data is the average of three replicates and error is the standard deviation of the mean. F) Maximum likelihood tree of representative NAD-dependent DNA ligase proteins from BLAST hits to MIT9313 ND-Lig fl and MIT9312 ND-Lig tr. All sequences were classified as full-length or truncated based on polypeptide length. All truncated sequences were less than 260 residues and included only Ia and NT domains. Sequences were trimmed to remove non-aligned portions of full-length ligases prior to tree building.

## Results and Discussion

Our comparative analyses of 41 *P. marinus* genomes (13 complete, 28 scaffolds) found that in high-light strains of *P. marinus*, the genes for the ‘essential’ replicative ND-ligases are severely truncated encoding polypeptides of <250 amino acids (Fig 1B). These truncated ND-ligases (hereafter ND-Lig tr) lack the oligonucleotide-binding domain and DNA-binding elements (BRCT and Zn-finger) necessary for activity (6). All high-light *P. marinus* strains harbor at least two AD-ligases (AD-Lig P and AD-Lig B) and MIT9312 has a third AD-ligase AD-Lig W (Fig C). In contrast, low-light strains possess full-length ND-ligases with all domains required for DNA ligase activity (ND-Lig fl). No low-light strains have an AD-ligase, however members of low-light clade LLI, which occupies an intermediate evolutionary position between ecotypes have both full-length and truncated forms of ND-ligase.

Our extensive attempts to recombinantly produce ND-lig tr were unsuccessful as the expressed protein was insoluble; however, given the lack of essential catalytic domains, it is highly unlikely that these truncated proteins are active. The lack of an intact ND-ligase gene indicates that in the high-light strains, all replicative and repair functions must be carried out by one or more of its AD-ligases. To determine where these AD-ligases are located the high-light *P. marinus* genome, we carried out whole-genome alignments of representatives from each phylogenetic clade. Strikingly, we found that in high-light strains, AD-Lig P, occupies an identical genomic position to the replicative ND-Lig fl of the low-light ecotypes (Fig 1D). In all cases, the DNA ligase (either ND-lig fl or AD-lig P) is sandwiched between two small conserved hypothetical proteins and downstream of the replicative recombinase RecA. This analogous sequence context indicates that AD-Lig P has genetically substituted the ND-ligase in high-light strains, and suggests it may also functionally replace it in DNA replication. As we have previously described, AD-Lig P is active on singly nicked or cohesive breaks and prefers Mg as a divalent cation (4, 7). We further confirmed the predicted cofactor preferences of ND-Lig fl and AD-Lig P as NAD and ATP by assaying purified recombinant protein (Fig 1E). We also demonstrated through comparison of specific activities, that AD-Lig P is more than 10 fold as effective at sealing singly-nicked double-stranded DNA substrates relative to the low-light ND-Lig fl under equivalent conditions.

Genome rearrangements have played a key role in the evolution of *P. marinus* ecotypes (8), therefore we investigated whether any of these DNA ligases are located within mobile regions. No bacteriophages or genomic islands are predicted in regions containing ND-Lig fl, AD-Lig P, or AD-Lig B, however AD-Lig W which is only found in only a few strains is located in a known high-light genomic island (9). The truncated ND-ligases are also located in these genomic islands and their positions in *P. marinus* genomes vary considerably. No synteny in genes adjacent to ND-lig tr was found between different high-light strains, or with genes flanking the full-length ND-ligases of low-light ecotypes. In addition, both ND-lig tr genes of low-light LLI strain NATL2A (which also has a full-length ND-ligase) are located in an incomplete prophage, suggesting that these truncated ligases are not simply pseudogenes of a former replicative ligase, but may be horizontally transferred between strains. A phylogenetic tree of the ND-ligases supports this idea, showing that the truncated isoform forms a separate and more heterogenous clade (Fig 1F). Based on this, we propose that the ND ligases were truncated and duplicated in plastic regions of a sub-set of low-light strains, prior to replacement of the functional full-length ND-ligase by an ATP-dependent form in high-light strains.

## Conclusions

To summarize, our genetic and biochemical analyses show that: 1) All high-light strains of *P. marinus* lack a gene encoding a complete ND-ligase gene, meaning this essential role must be fulfilled by one of its AD-ligases; 2) The AD-ligase AD-Lig P is substituted in the equivalent genetic context of the ND-ligase suggesting this isoform is responsible for replicative ligase activity; 3) The high-light AD-Lig P is a *bona fide* ATP-dependent DNA ligase with >10x higher specific activity relative to the low-light ND-ligase. The finding that an ATP-ligase must carry out both repair and replication in high-light *P. marinus* overturns the present paradigm of a clear evolutionary split between AD- and ND-ligase usage. This evidence that high-light strains of *P. marinus* are an evolutionary exception to the rules of replication recommends the various ecotypes as model systems to address the question of why most bacteria exclusively employ ND-ligases for DNA replication. As advances are being made towards genetic manipulation of *P. marinus* it may soon be possible to address this aspect directly (10).

## Materials and Methods

Complete genomes for *P. marinus* were downloaded from ProPortal and NCBI (11) (see S1 for strain details and accession numbers). DNA ligases were identified in all genomes by searching for the nucleotidyltransferase domain of ND-ligases (PF01653) and AD-ligases (PF01068). Fulllength coding sequences were retrieved and additional ligase-associated domains detected using Pfam as described previously (4). Whole genome alignments were constructed using progressiveMAUVE with default settings (12). Bacteriophage and genomic islands were identified using Alien_hunter (13) and Phaster (14). Phylogenetic trees used the Maximum likelihood method with Tamura-Nei model (DNA gyrase B) or using the JTT model (ND-ligases) and 1000 bootstrap replicates. Recombinant ND-Lig fl (MIT9211, WP_012196358) was expressed and purified as described for AD-Lig P (7), and both proteins were assayed described therein.

## Acknowledgments

We thank the Marsden Fund of New Zealand (18-UOW-034, AW) and Research Council Norway (244247, AW and EH) for funding. ERS is supported by a University of Waikato study award.

**Supplementary materials for this manuscript include the following:**

Datasets S1 to S2

**Dataset S1.** DNA ligases detected in 41 complete and partial genomes of *P. marinus* strains

**Dataset S2.** Whole genome alignment of representative strains from P. marinus ecotypes MIT_9312 (HLII), MED4 (HLI), NATL2A (LLI), MIT_9211 (LLII/III), MIT_9313 (LLIV). Genomes were aligned in progressiveMauve using default settings (Darling AE et.al (2010) *PLoS One* 5(6):e11147).

